# Identification of the associated expression patterns as potential predictive markers for cancer prognosis

**DOI:** 10.1101/2020.03.30.015180

**Authors:** Han Liang, Cong Lin, Yong Hou, Fuqiang Li, Kui Wu

## Abstract

Dysregulated gene expression can develop as a consequence of uncontrolled alterations of tumor cells. Analysis of these abnormal alterations will improve our understanding of the tumor development and reveal the corresponding clinical associations. It is well known that multiple genetic abnormalities could be observed in the same tumor, however, the interactions between those abnormal events are rarely analyzed. To address this problem, we constructed a novel gene expression correlation network by integrating the transcriptomes of 5,001 cancer patients from 22 cancer types. We investigated how the change of associated expression pattern (AEP), which describe certain associations between gene expression, could affect the cancer patient’s prognosis. Consequently, we identified an AEP composed of mitosis-related gene expressions, which is significantly correlated with overall survival in most cancer types. In particular, the AEPs could present the association between gene expressions and show distinct effects on prognosis prediction for cancer patients, suggesting that AEP analysis is indispensable to uncover the complex interactions of abnormal gene expressions in tumor development.

## Main

Dysregulated gene expression in cancer cells would lead to disorders in different cellular activities, including uncontrolled replication, proliferation and migration^1^. For example, overexpression of metallothionein 1E (MT1E) and nicotinamide N-methyltransferase (NNMT) could accelerate migration of bladder cancer cells and alter the expression of cell cycle related genes which will in turn to affect cancer cell proliferation^2,3^. Studying those abnormal expression events could improve our understanding of the role genes play in tumor’s development, and reveal potential applications of gene expression in facilitating the development of cancer treatment. Compelling evidence indicates that abnormal gene expression associates with patient’s prognosis across different cancer types ^2–6^, however, the interactions between those abnormalities are poorly understood. These interactions may influence the way we interpret the function of those events in tumor development and patient’s prognosis prediction. Hence, a clear understanding of relationships among gene expressions is essential.

To analyze the relationships among genes and to understand how the alteration of these relationships could change the way gene affects patient’s prognosis, we designed series analysis based on a novel gene expression correlation network. As human genome has tens of thousands of genes, it’s inefficient to analyze the relationship among all genes, which would also lead to an unacceptable high percentage of false positives. To overcome this problem, we only focused on the genes which display more connections with others in the expression network. We divided those genes into subsets according to their predictive effect on patient’s prognosis and analyzed their relationships within subsets. The genes from the same subset showed significant correlations with each other, and the associated expression patterns (AEPs) formed by them might actually be vary between groups with different characteristics of patients. In the present study, we demonstrated the strong influence of AEPs on predictive effect of patient’s prognosis. Our results suggested that AEPs could be used as a powerful tool to explore complex gene-gene interactions during tumor development.

We performed our analysis on transcriptomic data from TCGA ^8^. In the data arrangement section (Fig.1), we filtered patients according to their clinical information. We used the progression-free interval (PFI) time in the following analysis to define patient’s survival situation, as PFI is generally considered to be the better clinical endpoint choices than over survival (OS) and disease-specific survival (DSS) ^9^. We removed all patients without the PFI value and removed cancer types which patient sizes were less than 40. In total, 5,001 patients from 22 cancer types were included. Next, we removed the genes with low-quality expression data and eventually retained 11,241 common genes of all 22 cancer types. Specifically, we considered a gene’s expression data to be low quality when its expression values were the same for more than 5% patients in a single type of cancer, as the repeated values were generated by the invalid 0s of reads per kilobase million (RPKM), which meant no reads covering the gene during sequencing.

**Fig.1.**
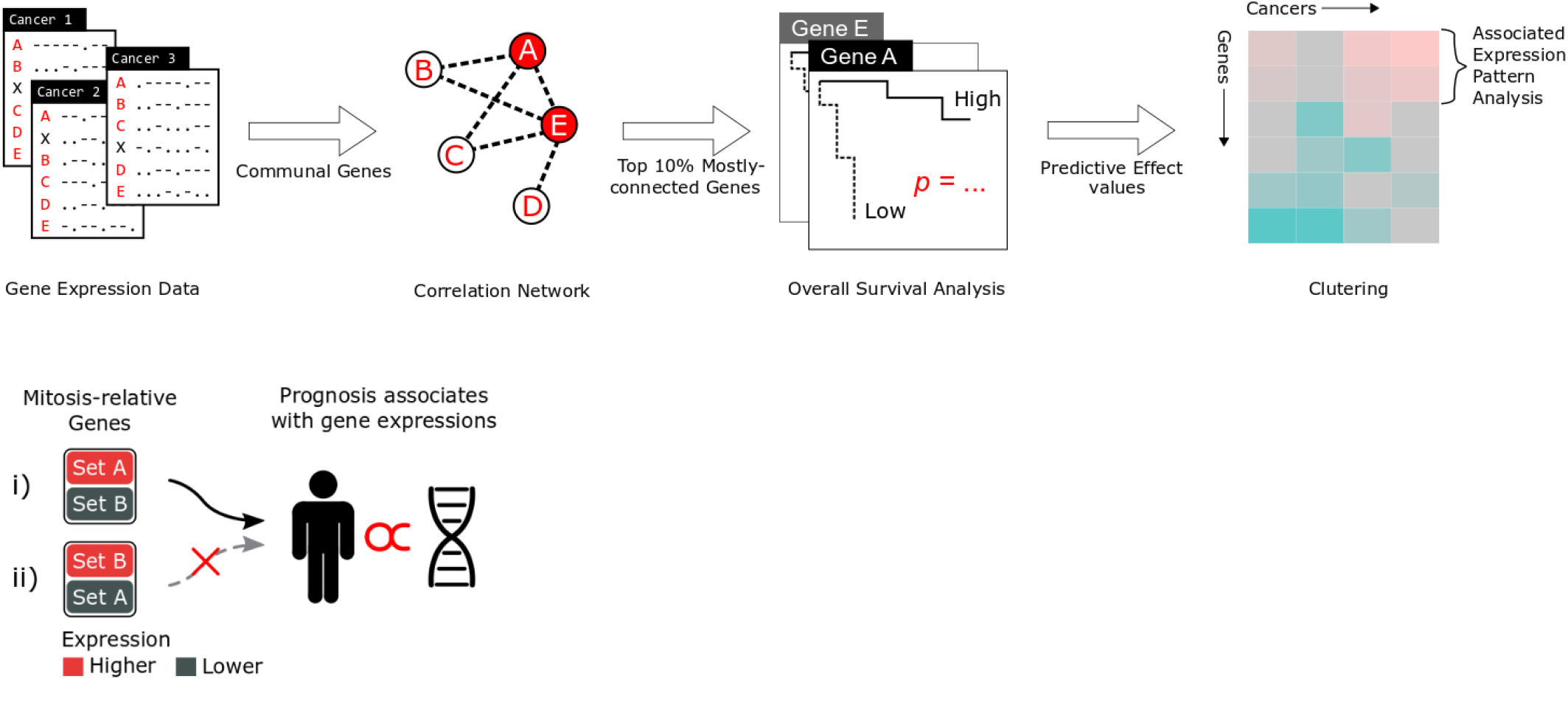
Schematic of the approach used to generate prognosis-relevant AEPs. The major process of our analysis could be divided into 4 major sections: 1) Data arrangement. Off-specification patients and low-quality data of gene expression was removed, while common genes of all cancer types were selected; 2) The correlation network construction. Genes were connected to each other with their direct or indirect correlations based on expression analysis; 3) Overall Survival (OS) Analysis. Ten percent of the most connected genes in the network were chosen, and the predictive effect of their expression on prognosis was assessed by Chi-square *p* values of the OS Analysis ^7^; 4) Clustering and analysis. Genes and cancer types with predictive values were clustered and AEPs of genes which formed the distant subset were analyzed; 5) AEPs verification. We found an AEP, the mitosis-relative AEP, and confirmed that when i) mitosis-relative genes’ expressions matched the AEP (clustered expression of Set A genes was higher than that of Set B), patients’ prognosis would likely associate with those gene expressions, otherwise ii) patients’ prognosis would unlikely associate with those expressions.

We next constructed a gene expression correlation network with retained data from TCGA, in which two genes would be connected if their expression levels were correlated with each other (the direct correlation) or with the third gene (the indirect correlation). Here, we made use of the indirect correlation to amplify aggregation effect of genes in the network. We then selected 10% of the genes that showed most connections with each other in the network, resulting in 1,125 genes for further analysis. This selection was based on our hypothesis that if a gene is associated with many other genes, it would be more likely to locate on a crucial position in the regulatory network. Subsequently, we assessed the predictive effect of these gene expressions on patients’ prognosis. We sorted patients in each cancer into the lower expression group and the higher expression group based on each gene’s expression, and measured the PFI difference between the two groups to assess predictive value of the gene expression on prognosis. Specifically, we measured the PFI difference between the two groups several times by taking different proportions of the groups each time, starting from 20% whilst increasing it by 5% until it reaches 50%. Therefore, we performed the OS Analysis^7^ 7 times on the different sizes of patients from the two groups, and got p1~p7, Chi-square *p* values which reflected the PFI difference between the two groups for each time. We then calculated the predictive effect value from p1~p7 with the formula (Fig.2a). The smaller *p* value (reflecting the more obvious PFI difference) means a stronger effect. If two groups show significant difference in prediction of PFI besides the significant difference in expression, it means that the gene expression associates with PFI. We used this method to calculated value for the predictive effect of each gene for all 22 cancer types, and then we clustered hub genes in the expression correlation network, which led us to a matrix of predictive value with rows as genes and columns as cancer types (Fig.2b).

**Fig.2.**
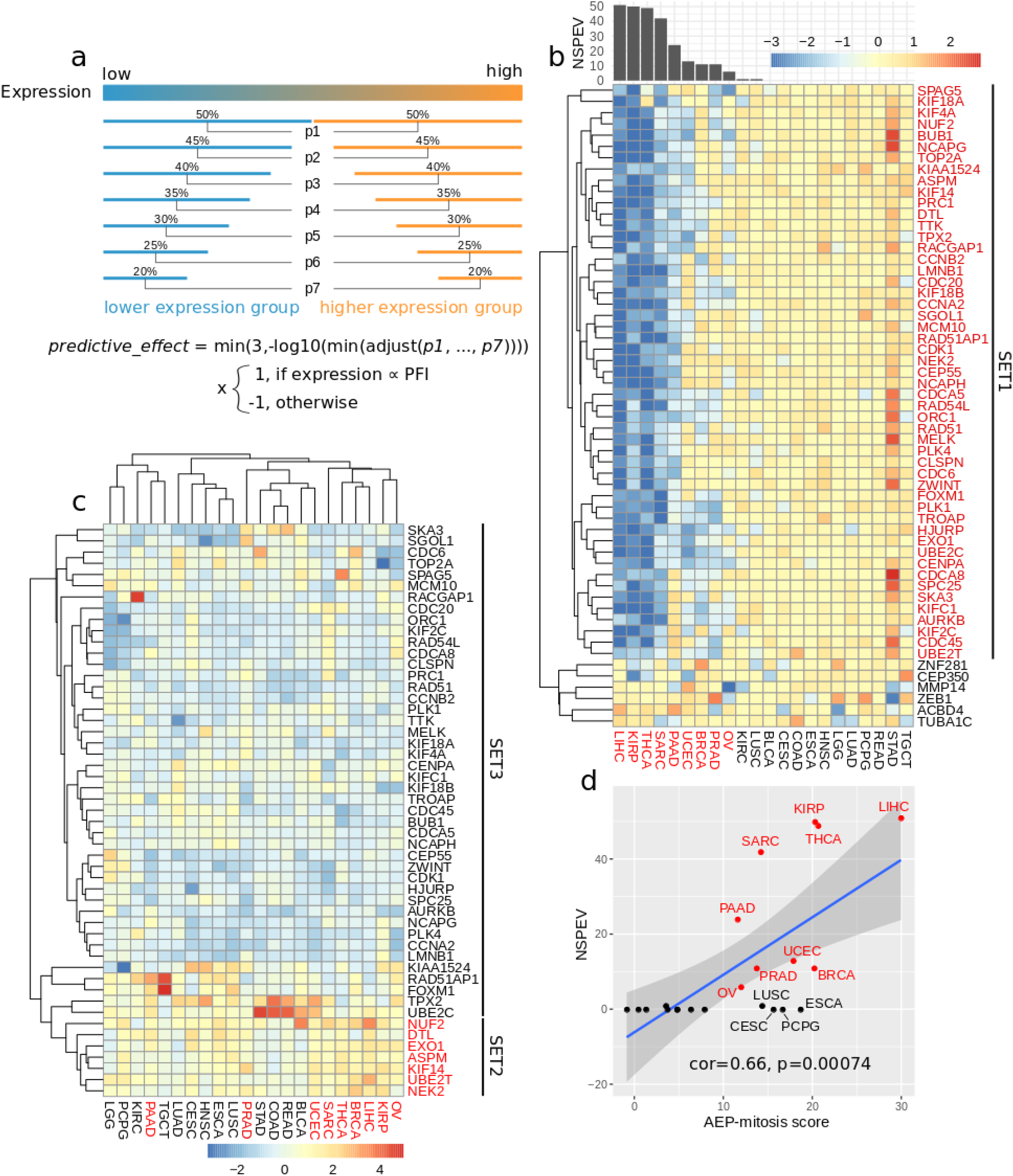
The discovery of a special AEP-mitosis for mitosis sensitive cancer types. **a) The method and formula used to calculate the predictive effect of each gene**. We calculated the PFI between two groups with significant difference of gene expression, and used a converted *p* value of the OS Analysis to measure the predictive effect. The function “*adjust*” used in the formula calculated the False Discovery Rate (FDR) of p1~p7, which returned the same number of adjusted *p* values. To avoid patient’s overlap between the two crowds, we limited the percentage no more than the 50% (the half), and to avoid shortage of observations, we limited the percentage no less than 20%. **b) The clustered heatmap of genes and cancer types based on predictive effects**. The columns of the heatmap represent all 22 cancer types and the rows represent part of genes with significant predictive effect value (the absolute value was larger than −log(0.05)). To illustrate it properly, all predictive effect values are limited to locate in the range of [−3, 3], any values that larger than the 3 (or smaller than −3) have been set as 3 (or −3). Under-zero values (bars in blue) represent that lower gene expression associates with better prognosis (LEBP), and above-zero values (bars in red) represent that higher gene expression associates with better prognosis (HEBP). Genes in red are referred as SET1 (including 51 genes) in the following texts. Most of mitosis-related genes (SET1, for the type of LEBP) with higher predictive effect values (the absolute value ≥ 10) was frequently found in cancer types marked in red. The cancer types were sorted according to their NSPEVs (represented by the barplot above). **c) Normalized gene expression landscape of SET1 across different cancer types**. The expression value has been normalized in each individual cancer separately, which means that values couldn’t be compared across cancer types. Genes in red are referred as SET2, and the rest genes in black are referred as SET3. The cancer types in red will be referred as mitosis-sensitive cancer types in the following texts. **d) The correlation between NSPEV and AEP-mitosis score in each individual cancer.** The correlation value and the *p* value are calculated with the Pearson’s product moment correlation method. AEP-mitosis score was calculated by subtracting the sum of SET2 gene expression with that of SET3.

From the clustered results, we discovered a distinct gene subset (Fig.2b, SET1). Genes in SET1 have similar values of predictive effect across cancer types and are mainly related to mitosis, including kinesin-like protein family series genes (KIFC1, KIF18B, KIF4A, KIF14 and KIF18A) ^10^, cell division cycle genes (CDC20, CDCA5, CDC6, CDC45 and CDC48)^11^ and some other genes (such as UBE2C^12^, CDK1^13^, BUB1^13^), which are all crucial for cell reproduction and tumor development^14^. Interestingly, these genes show more impact on certain cancer types, like KIRP, LIHC, SARC, THCA, OV, PAAD, UCEC and BRCA (mitosis-sensitive cancer). To assess whether this specific effect is due to variety of expression of those mitosis-related genes, we studied these genes’ expression patterns and associations between them. We tried to figure out whether the change of SET1 gene’s AEP could lead to the corresponding change in predictive result.

We first found a special AEP for most mitosis-sensitive cancer types from the expression landscape of SET1 genes (Fig.2c). For those mitosis-sensitive cancer types, the expressions of genes like NUF2,DTL,EXO1,ASPM,KIF14,UBE2T and NEK2 (genes in red in Fig.2c, referred as SET2) are higher than that of the rest genes (genes in black in Fig.2c, referred as SET3), which forms a distinct AEP (AEP-mitosis) from other non-mitosis-sensitive cancer types. We next verified this AEP-mitosis by examining the correlation between the number of genes with significant predictive effect values (NSPEV) and the expression difference between SET2 and SET3 in each individual cancer (Fig.2d). The Pearson’s product-moment correlation of them is ≈0.66 with a *p* value≈0.00074, which shows the expression difference associates with the NSPEV positively. For all mitosis-sensitive cancer types, the AEP-mitosis scores for expression differences between SET2 and SET3 are higher than 10, it thus seems that AEP of SET2 and SET3 is specific for this type of cancer. However, there are four non-mitosis-sensitive cancer types (LUSC, CESC, PCPG and ESCA) also with high AEP-mitosis score (Fig.2d). Among these four cancer types, three (LUSC, ESCA and CESC) are cancer types with squamous morphology components in Hoadley et al. study^15^ and all 5 cancer types of this kind (BLCA, CESC, ESCA, HNSC and LUSC) are non-mitosis-sensitive cancer types in our result, which suggests cancer types with squamous morphology might not follow the AEP-mitosis.

To corroborate AEP-mitosis could influence predictive effects of mitosis-related genes, we perform analysis from the gender aspect. Patients from 16 types of cancer were divided by genders in the following analysis, including BLCA, COAD, ESCA, HNSC, KIRC, KIRP, LGG, LIHC, LUAD, LUSC, PAAD, PCPG, READ, SARC, STAD and THCA (except for the gender-related cancer types, such as BRCA, OV, PRAD, CESC). Of those cancer types, 5 cancer types (KIRP, LIHC, LUAD, SARC and THCA) showed more than 3 NSPEVs difference between two gender groups (referred as gender-sensitive cancer types; Fig.3a and NSPEV in Tab.1). We found that for these 5 cancer types, a group with a higher AEP-mitosis score is always accompanied with a higher NSPEV, compared to the other group from the same cancer indicating that the change of AEP-mitosis led to the change of predictive effect of genes. We also confirmed that the AEP-mitosis could influence the predictive effect of the clustered expression (CE) of SET1. We used the median value of CE, since distribution of CE is skewed and median value can truly reflect the middle of the data set. We selected patients from the highest (or the lowest) 30% SET1-CE to be the higher-SET1-CE (or the lower-SET1-CE) crowd in a single gender group and checked the PFI difference between the two crowds. We found that a gender group with a higher AEP-mitosis score always shows a smaller *p* value (reflecting a more significant difference in PFI between its two crowds) comparing to the other group in the same cancer (Fig3b and *p* value in Tab.1), suggesting that the predictive effect of SET1-CE on PFI is strengthened with the increase of AEP-mitosis score. As the male group and female group both came from the same cancer, it could exclude other possible clinical influences on the prediction of prognosis between these two groups. These results suggest AEP-mitosis effectively contribute to patient’s prognosis prediction based on the expression of mitosis-relative genes.

**Fig.3.**
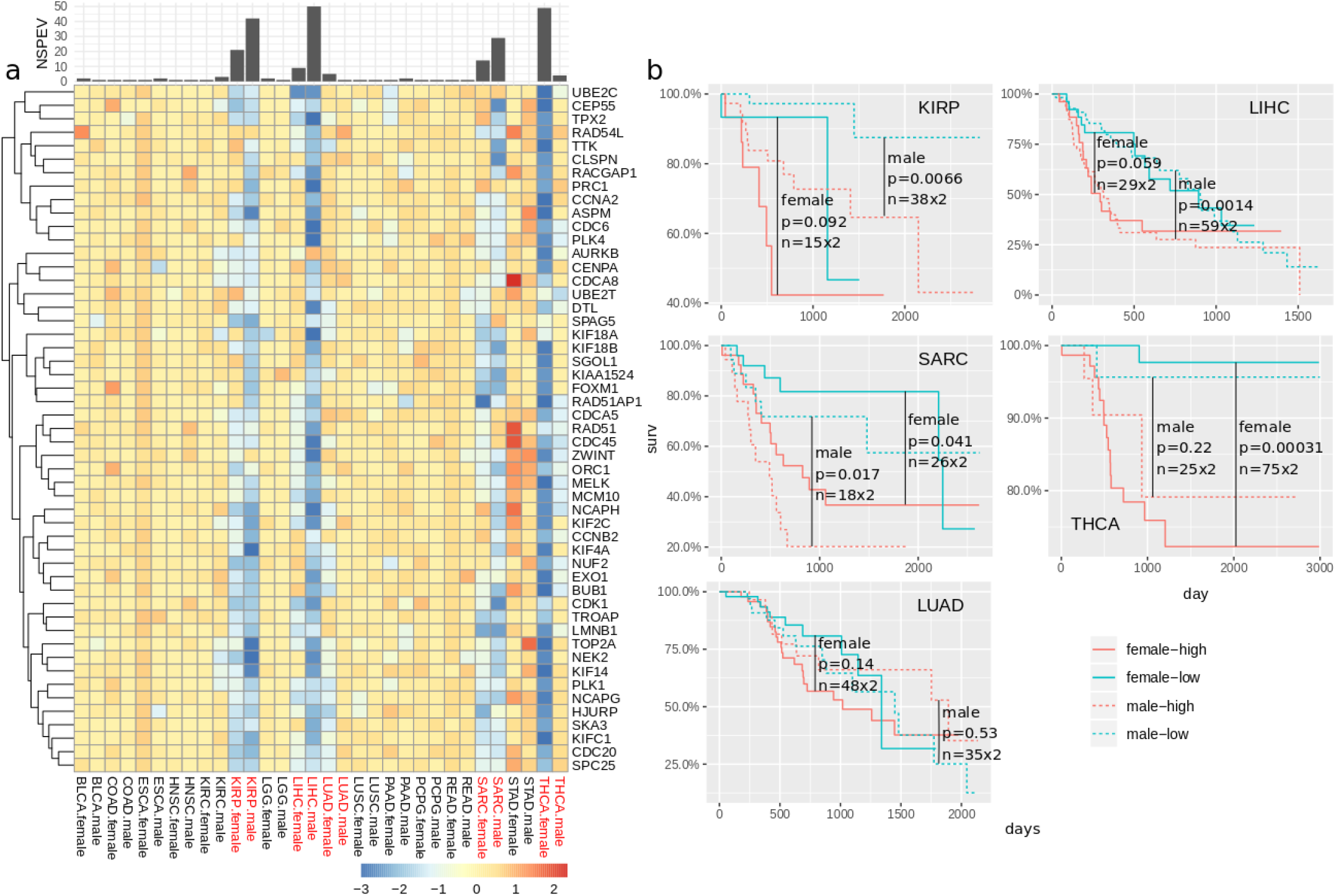
The AEP-mitosis could influence predictive effects of mitosis-related genes. a) Predictive effects of SET1 genes in different gender groups of 16 non-gender-related cancer types. b) The colors of bars have the same meanings as that of Fig.2b. Gender groups of the cancer marked in red show clear NSPEVs difference (>3) between each other (5 gender-sensitive cancer types in total). Results of the OS Analysis on the lower-SET1-CE crowd and the higher-SET1-CE crowd in 5 gender-sensitive cancer types.

**Tab.1.**
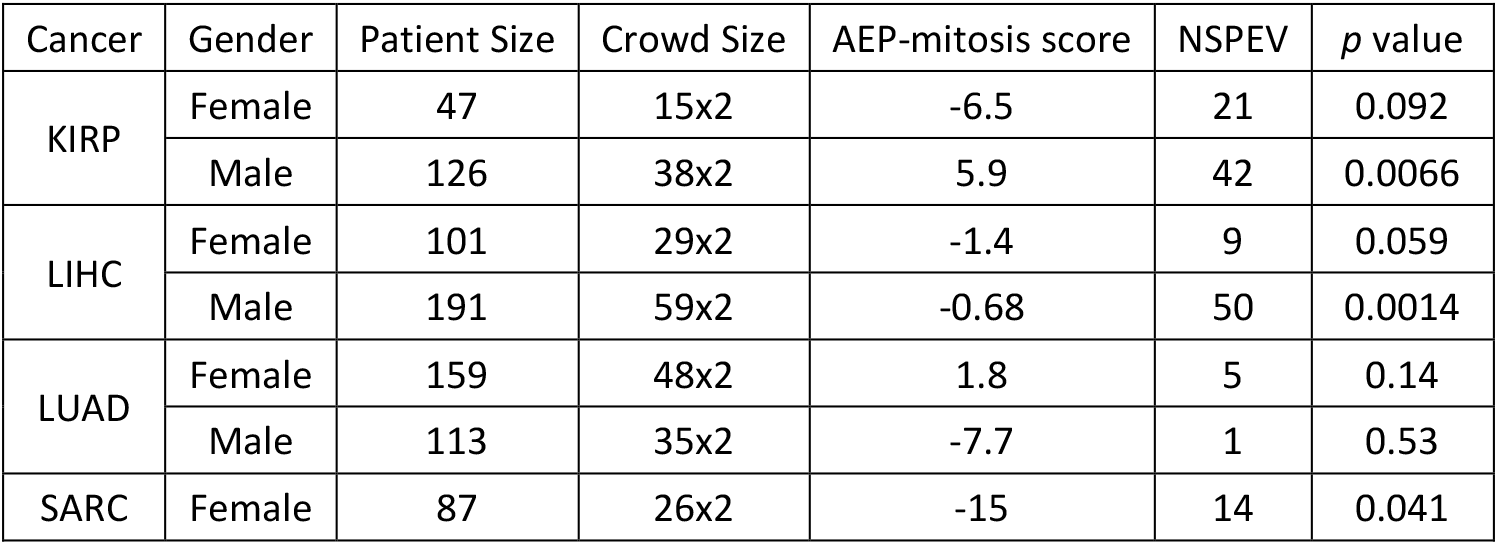
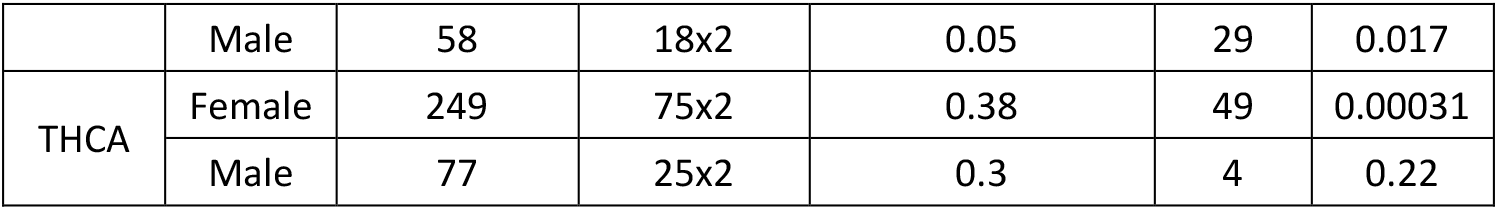
Statistic values of different gender groups of 5 gender-sensitive cancer types.

The Patient Size refers to the number of patients of each group. The Crowd Size refers to the number of patients we took to form the lower-SET1-CE crowd and the higher-SET1-CE crowd from the same group. The NSPEV refers to the number of genes with significant predictive effect value from SET1. The *p* value refers to the *p* value of the OS Analysis based on the lower-SET1-CE crowd and the higher-SET1-CE crowd.

Tumor variations between genders have been comprehensively studied^15^, yet the mechanism of different prognosis between genders in some cancer types remains elusive. Our results demonstrated different mitosis-related AEP between genders, which might provide a possible explanation for the gender-dependent prognosis. Importantly, the AEP-mitosis itself also associated with cancer patient’s PFI. We checked the PFI difference between two crowds with different range of AEP-mitosis score in each cancer. We selected patients with the highest 30% AEP-mitosis score as the HIGH crowd, and patients with the lowest 30% AEP-mitosis score as the LOW crowd and measured the PFI difference between the two crowds with *p* value of the OS Analysis (Tab.2). Of all 22 cancer types, 8 cancer types (BRCA, KIRP, LIHC, LUSC, PAAD, SARC, STAD and THCA) presented significant PFI difference (*p* value <=0.05) between the two crowds. Similar to the clustering result based on predictive effect (Fig.2b), that STAD was the only cancer type in which higher SET1 gene expression associates with better prognosis, STAD in this analysis again showed a different effect. In STAD, the LOW crowd owned the better PFI than the HIGH crowd, which was distinguishable from the other 7 cancer types. Notably, 6 of these 8 cancer types (expect for LUSC and STAD) were mitosis-sensitive cancer types, suggesting that AEP-mitosis could hardly affect non-mitosis-sensitive cancer types.

**Tab 2.**
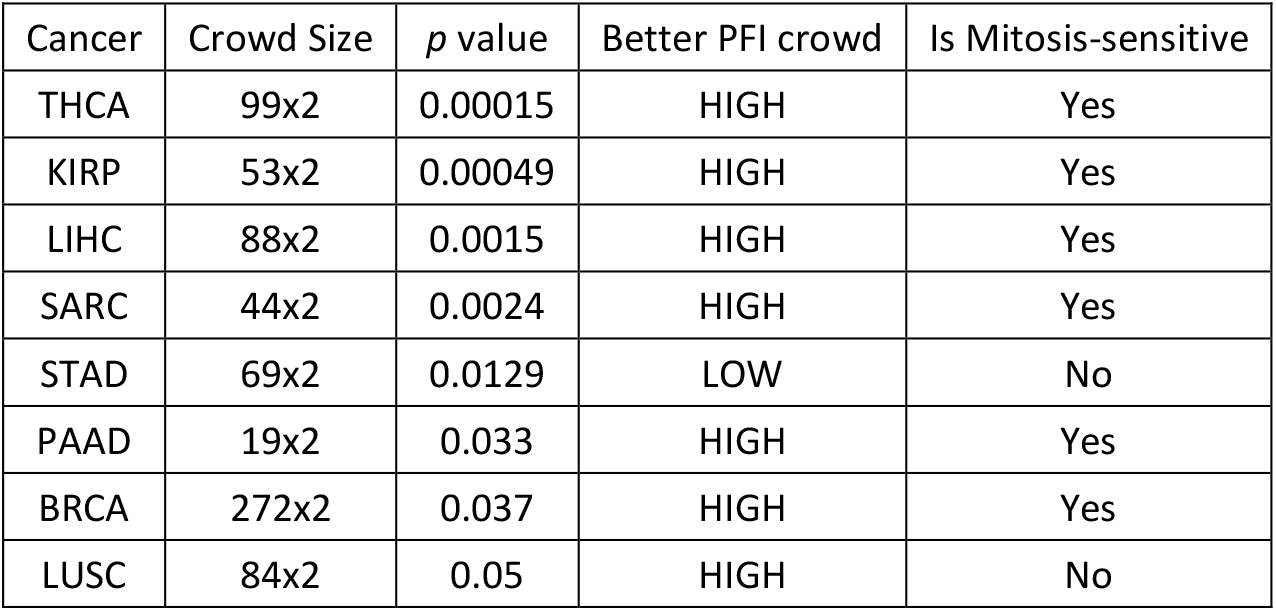
The PFI difference between crowds divided by AEP-mitosis scores. Patients were divided into two equal crowds based on the AEP-mitosis score in each cancer. Only cancer types with significant *p* value were listed here.

The overexpression of BUB1 and UBE2C(UbcH10) might be the reason for the different predictive effects of SET1 genes in STAD. BUB1 is a mitotic checkpoint gene, its overexpression would cause chromosome instability and further lead to poor prognosis for several cancer types^16,17^. However, as shown in Fig.2b, patients of STAD with higher expression of BUB1 show better prognosis^18^, which is distinct from the rest of SET1 genes. Intriguingly, UBE2C’s overexpression is also the most obvious feature of STAD (Fig.2c). UBE2C encodes a member of the E2 ubiquitin-conjugating enzyme family, which is required for the destruction of mitotic cyclins and for cell cycle progression ^12^. Given the precondition that UBE2C overexpression could cause chromosome mis-segregation^19^, BUB1 overexpression would aggravate chromosome instability, the two additive effects may accelerate tumor cell death ^18,20^, making BUB1 overexpress become a special benefit for STAD patients. It is noteworthy that three gastrointestinal cancer types, STAD, COAD and READ, exhibit abnormally high expression of UBE2C, suggesting that this gene may play a special role in intestinal tissues.

In the present study, we only analyzed limited amount genes from the gene expression correlation network, and there was still a large number of genes in the network left to be investigated. As the correlation network brought mitosis-related genes to our attention, we believe that this network is effective in revealing the crucial genes and will make a profound impact on the analysis of emerging multi-omics data.

SET2 and SET3 both come from SET1, when the AEP between SET2 and SET3 changes, the prognosis predictive effect of SET1 genes expression also changes accordingly(Fig3.b), which suggests that AEP could be used to understand the complex interactions between genes during tumor development and to reveal the extensive contribution of different gene-gene association to prognosis of cancer. Genes from the same pathway interact with each other through different ways, including assistance, substitute and competition, and the relative strength of the gene expression could determine the pathway’s effect. Inside a healthy individual body, the associations among genes could be in relative equilibrium states; but when the states are broken, it would cause unexpected consequences. There are still many unknowns about AEPs, i.e. the detailed molecular and cellular explanations of AEPs, which requires further exploration. We believed that AEPs could be an essential tool for cancer research in the further.

## Methods

### 1. Data Arrangement

We downloaded the level 3 data of gene expression (*.mRNAseq_Preprocess.Level_3.2016012800.0.0.tar.gz) and level 4 data of copy number (*.CopyNumber_Gistic2.Level_4.2016012800.0.0.tar.gz) from Broad GDC Firehose (https://gdac.broadinstitute.org), and normalized the Z-score result (*.uncv2.mRNAseq_RSEM_Z_Score.txt) with copy number result (all_thresholded.by_genes.txt) by CBioPortal convertExpressionZscores method (https://github.com/cBioPortal/cbioportal/blob/master/core/src/main/scripts/convertExpressionZscores.pl). We further filter the normalized gene expression data with the following strategies:

a. removed patients without platinum-free interval (PFI);
b. removed cancer types that owned less than 50 patients;
c. removed genes that more than 5% patients with the same expression value in a cancer type;
d. removed non-communal genes of all cancer types.

We finally kept 11241 genes and 5001 patients involving 22 cancer types, including BLCA(n=187), BRCA(n=953), CESC(n=174), COAD(n=188), ESCA(n=88), HNSC(n=130), KIRC(n=117), KIRP(n=184), LGG(n=136), LIHC(n=318), LUAD(n=302), LUSC(n=300), OV(n=149), PAAD(n=69), PCPG(n=163), PRAD(n=338), READ(n=47), SARC(n=155), STAD(n=241), TGCT(n=110), THCA(n=357), UCEC(n=424).

### 2. Correlation Networks Construction

a) The times connected by other genes of each gene in the gene expression correlation network was counted by the following algorithm:

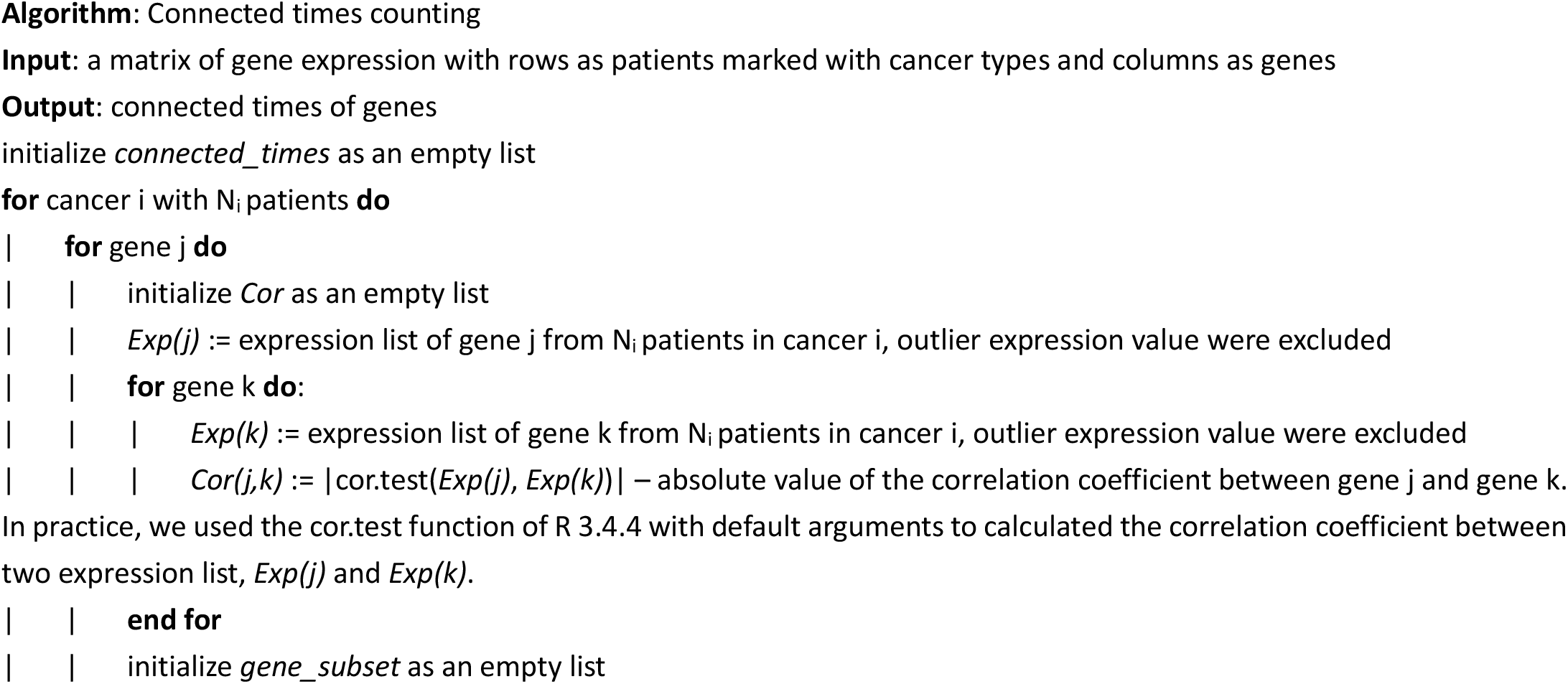
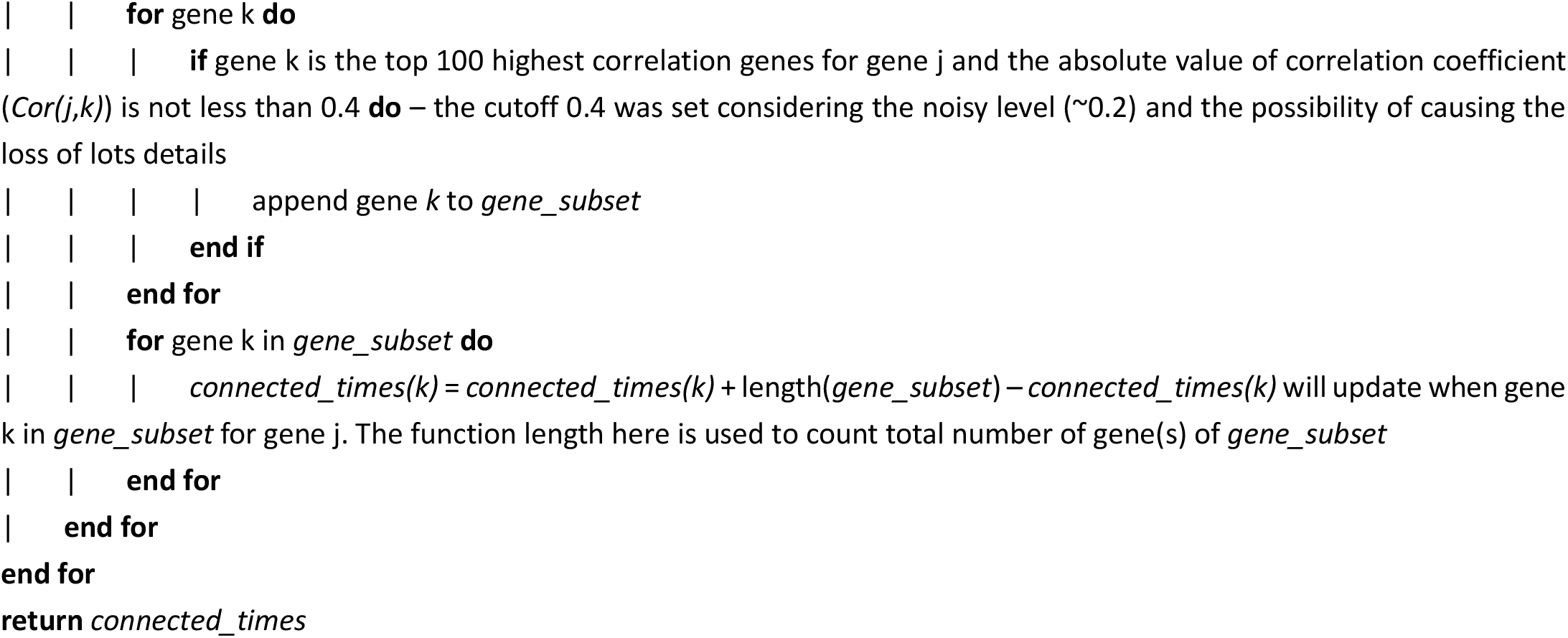

Notice the step to update *connected_times* “*connected_times(k)* = *connected_times(k)* + length(*gene_subset*)”, since all the genes in *gene_subset* both show obvious correlation with the gene j in preceding text, it means that those genes correlate with each other indirectly. As we allowed indirect method to amplify aggregation effect of genes in the network, the connected times of those genes will increase by the number of those genes. An expression value will be considered as an outlier one when its value is beyond the range [quantile_first_−1.5*( quantile_third_− quantile_first_), quantile_third_+1.5*( quantile_third_−quantile_first_)] ^21^.

b) 1125 genes with the highest 10% connected times were selected for following prognosis analysis.

### 3. Prognosis analysis of gene expression

To assess the predictive effects of gene expression of highly connected genes in cancer patient, we use platinum-free interval (PFI) to perform survival analysis between *k*% patients with higher gene expression and k% patients with lower gene expression.

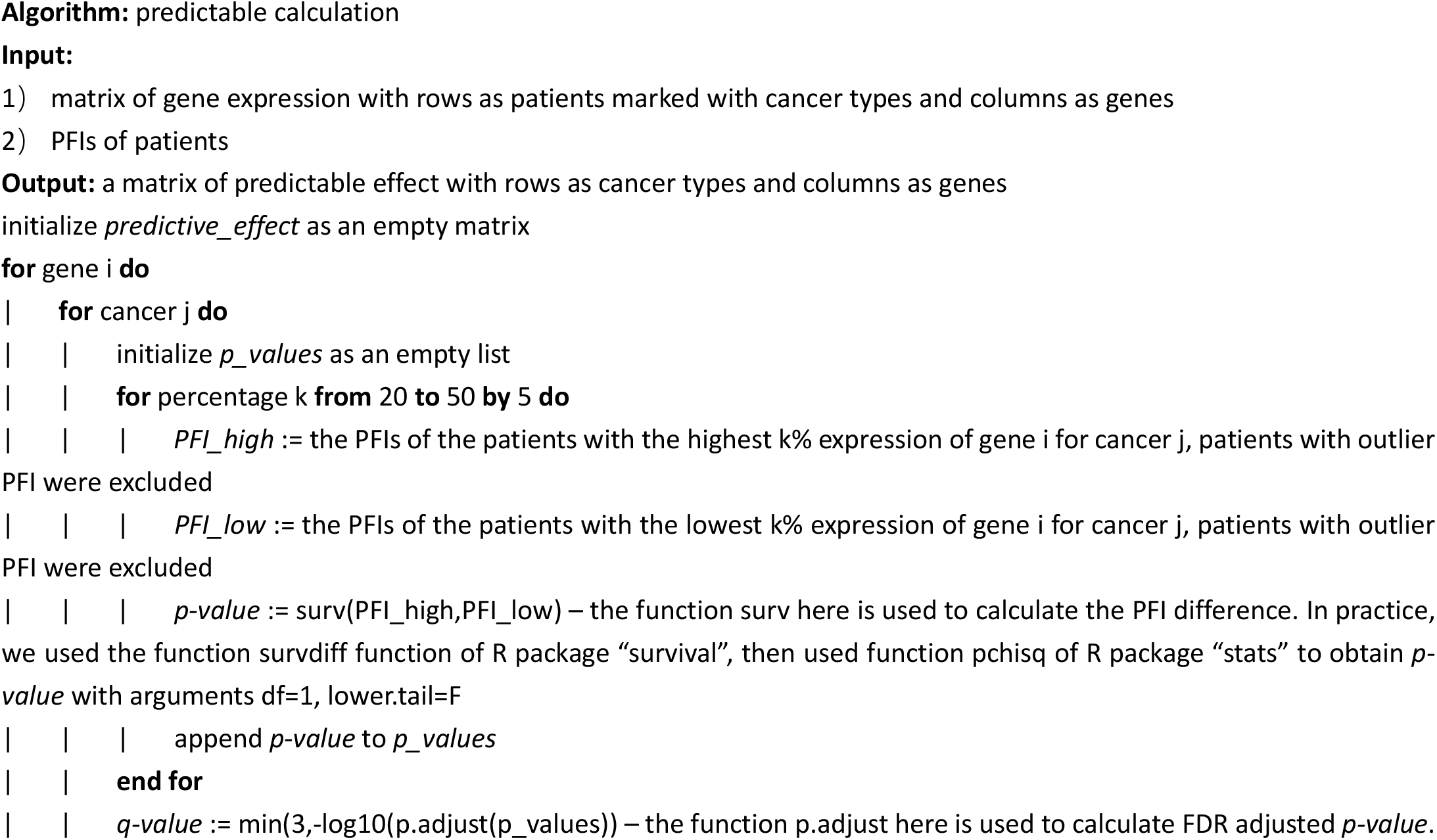
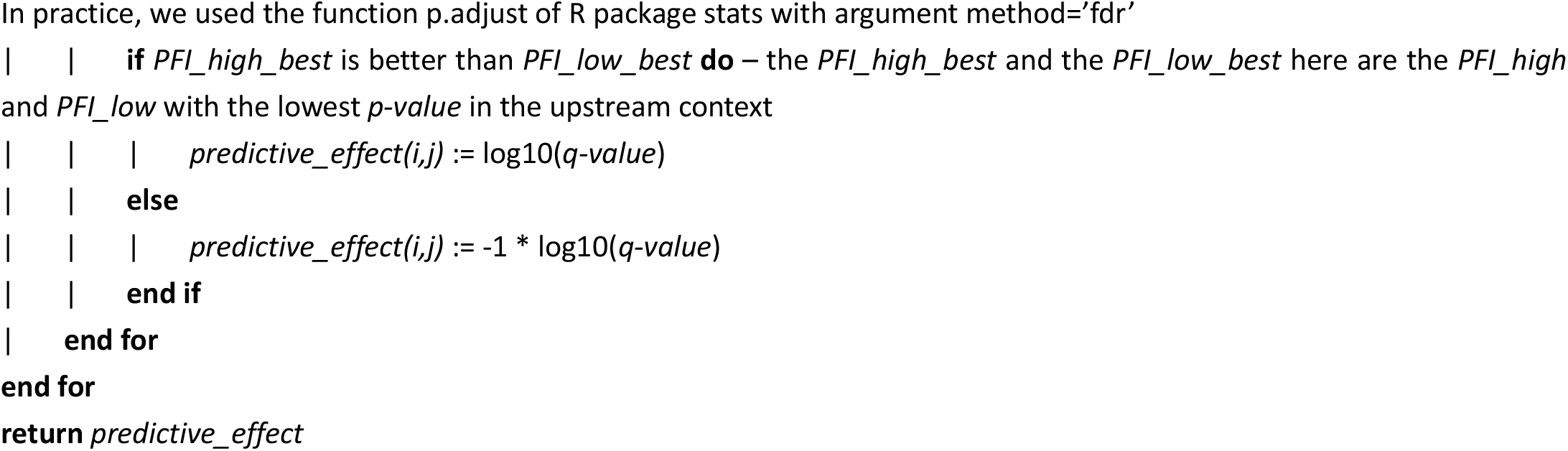

An PFI value will be considered as an outlier one when its value is beyond the range [quantile_first_−1.5*( quantile_third_−quantile_first_), quantile_third_+1.5*( quantile_third_− quantile_first_)] ^21^.

### 4. Associated expression patterns

a) Genes and cancer types were clustered by *predictive_effect* values (Fig.2b). Total of 57 genes with absolute *predictive_effect* value larger than −log(0.05) in at least 5 cancer types were kept. A branch of clustered tree with 51 genes (SET1) was related with mitosis.

b) Genes and cancer types were clustered by normalized gene expression value. The expression values were normalized with individual cancer type separately, which means that values couldn’t be compared across cancer types. Genes in red are referred as SET2, and the rest genes in black are referred as SET3. The cancer types in red will be referred as mitosis-sensitive cancer types in the following texts.

Notably, the AEP-mitosis scores in Tab.1 were calculated based on raw expression values instead of the normalized ones used in Fig2.d, as the raw expression values could be compared directly within a single cancer, which doesn’t require normalization.

## Author contributions

H. L. and C. L. conceived of the idea and wrote the manuscript. H.L. developed the model for Identification of the associated expression patterns. C. L. and F. Q. L. contributed to drafting and revising the manuscript. K.W., F. Q. L. and Y. H. supervised the work.

## Data and materials availability

We are thankful that The Cancer Genome Atlas (TCGA) and Broad Institute GDAC Firehose for sharing the data. Level 3 data of gene expression (*.mRNAseq_Preprocess.Level_3.2016012800.0.0.tar.gz) and level 4 data of copy number (*.CopyNumber_Gistic2.Level_4.2016012800.0.0.tar.gz) from Broad GDAC Firehose (https://gdac.broadinstitute.org).

We are thankful that Yan Liang (BGI-Shenzhen) for her meticulous project management supports.

## References

1. Bradner, J. E., Hnisz, D. & Young, R. A. Transcriptional Addiction in Cancer. Cell 168, 629–643 (2017).

2. Peng, X. et al. Molecular Characterization and Clinical Relevance of Metabolic Expression Subtypes in Human Cancers. Cell Rep. 23, 255–269.e4 (2018).

3. Knijnenburg, T. A. et al. Genomic and Molecular Landscape of DNA Damage Repair Deficiency across The Cancer Genome Atlas. Cell Rep. 23, 239–254.e6 (2018).

4. Wang, Y. et al. Comprehensive Molecular Characterization of the Hippo Signaling Pathway in Cancer. Cell Rep. 25, 1304–1317.e5 (2018).

5. Way, G. P. et al. Machine Learning Detects Pan-cancer Ras Pathway Activation in The Cancer Genome Atlas. Cell Rep. 23, 172–180.e3 (2018).

6. Schaub, F. X. et al. Pan-cancer Alterations of the MYC Oncogene and Its Proximal Network across the Cancer Genome Atlas. Cell Syst. 6, 282–300.e2 (2018).

7. Therneau, T. M. Survival Analysis. RStudio Package ‘ survival ’. 1–161 (2019).

8. Weinstein, J. N. et al. The Cancer Genome Atlas Pan-Cancer analysis project. Nat. Methods 45, 1113–20 (2013).

9. Liu, J. et al. An Integrated TCGA Pan-Cancer Clinical Data Resource to Drive High-Quality Survival Outcome Analytics. Cell 173, 400–416.e11 (2018).

10. Vicente, J. J. & Wordeman, L. Mitosis, microtubule dynamics and the evolution of kinesins. Exp. Cell Res. 334, 61–69 (2015).

11. Moriya, H., Shimizu-Yoshida, Y. & Kitano, H. In vivo robustness analysis of cell division cycle genes in Saccharomyces cerevisiae. PLoS Genet. 2, 1034–1045 (2006).

12. Bajaj, S. et al. E2 ubiquitin-conjugating enzyme, UBE2C gene, is reciprocally regulated by wild-type and gain-of-function mutant p53. J. Biol. Chem. 291, 14231–14247 (2016).

13. Diril, M. K. et al. Cyclin-dependent kinase 1 (Cdk1) is essential for cell division and suppression of DNA re-replication but not for liver regeneration. Proc. Natl. Acad. Sci. U. S. A. 109, 3826–3831 (2012).

14. Cho, S. Y., Kim, S., Kim, G., Singh, P. & Kim, D. W. Integrative analysis of KIF4A, 9, 18A, and 23 and their clinical significance in low-grade glioma and glioblastoma. Sci. Rep. 9, 1–14 (2019).

15. Hoadley, K. A. et al. Cell-of-Origin Patterns Dominate the Molecular Classification of 10,000 Tumors from 33 Types of Cancer. Cell 173, 291−304.e6 (2018).

16. Han, J. Y. et al. Bub1 is required for maintaining cancer stem cells in breast cancer cell lines. Sci. Rep. 5, (2015).

17. Ricke, R. M., Jeganathan, K. B. & van Deursen, J. M. Bub1 overexpression induces aneuploidy and tumor formation through Aurora B kinase hyperactivation. J. Cell Biol. 193, 1049–1064 (2011).

18. Stahl, D. et al. Low BUB1 expression is an adverse prognostic marker in gastric adenocarcinoma. Oncotarget 8, 76329–76339 (2017).

19. Van Ree, J. H., Jeganathan, K. B., Malureanu, L. & Van Deursen, J. M. Overexpression of the E2 ubiquitin-conjugating enzyme UbcH10 causes chromosome missegregation and tumor formation. J. Cell Biol. 188, 83–100 (2010).

20. Jeganathan, K., Malureanu, L., Baker, D. J., Abraham, S. C. & Van Deursen, J. M. Bub1 mediates cell death in response to chromosome missegregation and acts to suppress spontaneous tumorigenesis. J. Cell Biol. 179, 255–267 (2007).

21. Hearst, M. A. Untangling text data mining. in Proceedings of the 37th annual meeting of the Association for Computational Linguistics on Computational Linguistics - Proceeding, 3–10 (Association for Computational Linguistics, 1999).

